# Population diversity of cassava mosaic begomoviruses increases over the course of serial vegetative propagation

**DOI:** 10.1101/2021.03.22.436436

**Authors:** Catherine D. Aimone, Erik Lavington, J. Steen Hoyer, David O. Deppong, Leigh Mickelson-Young, Alana Jacobson, George G. Kennedy, Ignazio Carbone, Linda Hanley-Bowdoin, Siobain Duffy

**Affiliations:** Department of Plant and Microbial Biology, North Carolina State University, Raleigh NC 27695 USA; Department of Ecology, Evolution, and Natural Resources, Rutgers University, New Brunswick, NJ 08901 USA; Department of Entomology and Plant Pathology, Auburn University, Auburn, AL 36849 USA; Center for Integrated Fungal Research, Department of Entomology and Plant Pathology, North Carolina State University, Raleigh NC 27695 USA

**Keywords:** cassava mosaic begomoviruses, viral diversity, vegetative propagation

## Abstract

Cassava mosaic disease (CMD) represents a serious threat to cassava, a major root crop for more than 300 million Africans. CMD is caused by single-stranded DNA begomoviruses that evolve rapidly, making it challenging to develop durable disease resistance. In addition to the evolutionary forces of mutation, recombination, and reassortment, factors such as climate, agriculture practices, and the presence of DNA satellites may impact viral diversity. To gain insight into the factors that alter and shape viral diversity *in planta*, we used high-throughput sequencing to characterize the accumulation of nucleotide diversity after inoculation of infectious clones corresponding to African cassava mosaic virus (ACMV) and East African cassava mosaic Cameroon virus (EACMCV) in the susceptible cassava landrace Kibandameno. We found that vegetative propagation had a significant effect on viral nucleotide diversity, while temperature and a satellite DNA did not have measurable impacts in our study. EACMCV diversity increased linearly with the number of vegetative propagation passages, while ACMV diversity increased for a time and then decreased in later passages. We observed a substitution bias toward C→T and G→A for mutations in the viral genomes consistent with field isolates. Non-coding regions excluding the promoter regions of genes showed the highest levels of nucleotide diversity for each genome component. Changes in the 5’ intergenic region of DNA-A resembled the sequence of the cognate DNA-B sequence. The majority of nucleotide changes in coding regions were non-synonymous, most with predicted deleterious effects on protein structure, indicative of relaxed selection pressure over 6 vegetative passages. Overall, these results underscore the importance of knowing how cropping practices affect viral evolution and disease progression.

## Introduction

Begomoviruses (genus *Begomovirus*, family *Geminiviridae*) are single-stranded DNA (ssDNA) viruses that cause serious diseases in many important crops worldwide (1). They are characterized by their double icosahedral particles and whitefly (*Bemisia tabaci* Genn) vectors (2, 3). Like other ssDNA viruses (4–9), begomoviruses have the potential to evolve rapidly because of their small genomes, large population sizes, short generation times, and high substitution rates (10, 11). Begomoviruses also readily undergo recombination (12), but mutation is the major driver of diversification of begomovirus populations (13, 14). The highly polymorphic nature of begomovirus populations can lead to rapid adaptation and increased virulence (15, 16). Plant virus evolution is also influenced by ecological conditions, such as agricultural practices and environmental stresses (17–20).

Cassava is a major root crop in Africa (21), whose production has been severely impacted by a rapidly evolving complex of 11 begomoviruses species, 9 of which are in Africa, which cause Cassava mosaic disease (CMD) (22). Cassava mosaic begomoviruses (CMBs) have bipartite genomes that consist of two circular DNAs designated as DNA-A and DNA-B (23). Both genome components, which together are ca. 5.5 Kb in size, display high nucleotide substitution rates of approximately 10^−3^ to 10^−4^ substitutions per site per year (10), which are similar to the rates reported for RNA viruses (24, 25). DNA-A is necessary for viral replication, transcription and encapsidation, while DNA-B is required for viral movement (2, 3). Both genome components contain divergent transcription units separated by a shared 5’ intergenic sequence or common region that contains the origin of replication and promoters for viral gene transcription (2, 3). The viral replication origin includes a hairpin structure that contains the nick site for rolling circle replication and iteron motifs that function as origin recognition sequences (26, 27). The origin motifs are conserved between cognate DNA-A and DNA-B components.

DNA-A encodes six proteins via overlapping genes, while DNA-B encodes two proteins on nonoverlapping genes (2, 3). (See Supplementary Figure 1 for diagrams of viral clones and major functions of the viral proteins). Genes specified on the complementary DNA strand are designated as “*C*”, while genes on the virion strand are indicated as “*V*”.) The replication-associated protein (Rep, *AC1*), which catalyzes the initiation and termination of rolling circle replication (28) and functions as a DNA helicase (29), is the only viral protein essential for replication (30). REn (*AC3*) greatly increases viral DNA accumulation by facilitating the recruitment of host DNA polymerases for viral replication (31). Viral ssDNA generated during rolling circle replication can be converted to dsDNA (32) and reenter the replication cycle or be packaged into virions composed of the coat protein (CP, *AV1*) (33). TrAP (*AC2*) is a transcription factor (34). TrAP, AC4, and AV2 counter host defenses by interfering with post-transcriptional gene silencing (PTGS) and transcriptional gene silencing (TGS) (for review see(35). ACMV DNA-A also includes an *AC5* ORF of unknown function that overlaps *AV1* (36, 37). DNA-B encodes two proteins essential for movement. The movement protein (MP, *BC1*) is necessary for viral transport through the plasmodesmata into adjacent cells and systemically through the plant (38, 39). The nuclear shuttle protein (NSP, *BV1*) is involved in viral DNA trafficking into and out the nucleus and across the cytoplasm to the cell periphery in coordination with MP (38–40).

CMBs often occur in mixed infections leading to synergy between the coinfecting viruses and increased symptom severity (12, 41). In the 1990s and 2000s, synergy between *African cassava mosaic virus* (ACMV) and a recombinant CMB contributed to a severe CMD pandemic that spread from Uganda to other sub-Saharan countries and devastated cassava production (12, 42). In response to the pandemic, many African farmers adopted cassava cultivars with the CMD2 locus, which confers resistance to CMBs (43, 44). Cassava plants displaying severe, atypical CMD symptoms were observed in some fields after the widespread adoption of CMD2-resistant cultivars (45). Subsequently, two DNAs, SEGS-1 and SEGS-2 (<u>s</u>equences <u>e</u>nhancing <u>g</u>eminivirus <u>s</u>ymptoms), were shown to produce similar symptoms when coinoculated with CMBs into both resistant and susceptible cassava cultivars (45). SEGS-2, which occurs as episomes in CMB-infected cassava, virions and whiteflies, is thought to be a novel satellite (46). SEGS-1 also forms episomes in CMB-infected cassava but is likely derived from a cassava genomic copy (45). The uniqueness of SEGS-1 and SEGS-2 raises questions about how they interact with other viral components and might impact begomovirus diversity.

Little is known about how vegetative propagation, environmental factors, and the presence of the SEGS affect CMB evolution. To generate knowledge about the impact of these factors, we examined viral diversity in mixed infections of ACMV and *East African cassava mosaic Cameroon virus* (EACMCV) during serial vegetative propagation of cassava plants inoculated with only CMBs or coinoculated with SEGS and grown under controlled conditions at two temperatures. This study provided evidence that vegetative propagation, but not temperature or the presence of SEGS, had a significant impact on viral diversity over time.

## Methods

### Vegetation Propagation Study with Two Passages (Veg2) at Two Temperatures

Cassava plants (*Manihot esculenta* cv. Kibandameno) were propagated at 28°C and 30°C under a 12-h light/dark cycle representing the predicted 2°C temperature shift in Africa by 2030 (47) and inoculated using a microsprayer to deliver plasmids (100 ng) containing partial tandem dimers of DNA-A or DNA-B of ACMV (Accession Numbers: MT858793.1 and MT858794.1) and EACMCV (Accession Numbers: MT856195 and MT856192) (48, 49). Three plants were coinoculated with ACMV and EACMCV clones for each temperature treatment with each plant representing a biological replicate. Samples (1 mg) were collected at 28 days post inoculation (dpi) from symptomatic tissue near the petiole of leaf 2 (relative to the plant apex), flash-frozen in liquid nitrogen, and stored at −80°C until analysis. At 56 dpi, a stem cutting with two nodes was generated from each biological replicate and transferred to fresh soil after treatment with rooting hormone (Garden Safe: TakeRoot Rooting Hormone). The stem cuttings were sampled at 28 days after propagation as described above and used for the next round of propagation at 56 days. Propagated plants originating from the same inoculated source plant represent a lineage. Leaf tissue collection and propagation continued for a total of two rounds at each temperature following the above protocol. Frozen leaf tissue was ground, total DNA was extracted using the MagMax^TM^ Plant DNA Isolation Kit (Thermo Fisher Scientific, Waltham, MA), and DNA < 6 Kb in size was selected using the Blue Pippin Prep system (Model # BDQ3010, Sage Science, Beverly MA) prior to library prep (50).

### Vegetative Propagation Study with Six Passages (Veg6)

Kibandameno plants were propagated from stem cuttings, grown at 28°C, and inoculated as described above. Plants were inoculated with plasmid DNA (100 ng) corresponding to ACMV + EACMCV, ACMV + EACMCV + SEGS-1 (AY836366), or ACMV + EACMCV + SEGS-2 (AY836367). Each treatment was replicated 3 times. Leaf samples were collected, and stem cuttings were propagated as described above. This process was repeated six times with a total of seven rounds including the initial inoculated plant. Total DNA was isolated but was not size-fractionated because earlier serial experiments showed that sequencing total DNA samples produce sufficient read coverage for viral diversity analysis (50).

### Library Preparation

Sequencing libraries for Veg2 and Veg6 were generated using EquiPhi polymerase (Thermo Fisher Scientific, Waltham MA) for rolling circle amplification (RCA) and the Nextera XT kit (Illumina, San Diego CA) with unique dual index sequences for library preparation (50). The protocol was modified to include two RCA reactions for each sample. Each RCA reaction was diluted to 5 ng/mL, and 2 μL of each reaction were combined. The combined RCA reactions were diluted to 0.2 ng/μL (1 ng total in 5 μL) and used to construct two technical replicate libraries. Libraries of the inoculum plasmids for ACMV and EACMCV (1 ng of plasmid DNA in 5 μL) were also prepared using the Nextera XT kit. The libraries were pooled in equimolar amounts and sequenced on an Illumina NovaSeq 6000 S4 lane to generate 150-bp, paired-end reads.

### Analysis of Raw Reads

Raw sequencing data were processed according to Aimone et al. (2020); the workflow is also available on Galaxy (ViralSeq, https://cassavavirusevolution.vcl.ncsu.edu/). Raw Illumina data is available from the NCBI Sequence Read Archive for Veg2 (PRJNA667210) and Veg6 (PRJNA658475).

### SNP filtering and functional analysis

SNPs were detected using VarScan (51) and filtered for all SNPs present in both of the sequencing technical replicates and at a frequency ≥3% in at least one replicate. SNPs present at ≥3% frequency in both technical replicates and present in more than one passage were selected for functional analyses. SNPs were categorized by intergenic region, protein, functional domain, and known motif using SNPeff (52). SNPs in coding regions were categorized as synonymous or non-synonymous. The SIFT tool D algorithm was used to predict the effects of single amino acid substitutions caused by non-synonymous codons, with a substitution scored as damaging (≤ 0.05) or tolerated (> 0.05) (53). Predictions are based on a scaled probability matrix built by the SIFT algorithm using protein sequence alignments between the queried protein sequence and proteins in known databases, in this case [non-redundant protein] (53). In the overlaps between coding regions, if a SNP was called using VarScan and identified by protein region using SNPeff in one coding region, its impact on the second coding region was categorized as synonymous or non-synonymous. SNPs in the common regions of the DNA-A and DNA-B of ACMV or EACMCV were compared to one another and the reference sequence of the infectious clone for each viral genome component used in the propagation studies. The alignments were performed using a Smith-Waterman sequence alignment (54) using SnapGene software (from Insightful Science; available at snapgene.com). A Mann-Whitney Test in R was used to calculate the difference between the frequency of SNPs observed in the historical database and SNPs that occurred in Veg6 experiment.

### Nucleotide diversity and Tajima’s D

Nucleotide diversity was calculated from SNP frequencies per nucleotide position by the formula *π* = ∑*_ij_ x_i_x_j_π_ij_* (55) using custom Python scripts. When calculating Tajima’s D, total read coverage averaged across all SNPs for a given region was used as a proxy for sample size (56).

Sliding window calculations of *π* were calculated by custom Python scripts with a window size of 300 bp and a step of 10 nucleotides and are reported at the central position of the window.

### Experimental effect analyses

The effect size of experimental design variables was analyzed in Python by linear regression using the statsmodels module (v0.12.0) with genomic nucleotide diversity as the response (57). Details of the model and results are discussed below.

### Analysis of nucleotide substitution bias

Filtered SNPs were combined by experiment, species, and component across all passages. Each unique substitution was used to generate observed counts. Substitution counts for each pair of nucleotides were tested by a *χ*^2^ test of independence on a 2X2 contingency table of observed and expected counts. To generate the expected counts for each pair, we assumed that substitutions in both directions were equally likely, divided observed counts in half, and adjusted for nucleotide proportions in the reference sequence (10). For example, considering the test for A↔T substitution bias in ACMV DNA-A in the Veg2 experiment, we observed a total of 5 A→T and 13 T→A substitutions. Counts of A and T in the ACMV DNA reference are 743 and 795, respectively, yielding expectations of 8.7 A→T and 9.3 T→A.

### Data and analysis availability

Data in the form of VarScan outputs for each of the models with associated metadata along with custom scripts used for analyses are available at https://www.github.com/elavington/PIRE.

## Results

### Experimental variable effects

We tested the effects of three treatments (temperature, vegetative propagation, and the presence of SEGS) on nucleotide diversity of ACMV and EACMCV in two experiments designated as Veg2 and Veg6. Veg2 tested the effect of temperature and included bombard plants (Passage 1, P1) and plants from two rounds of propagation (P2 and P3). Veg6 included clonally bombard plants (P1) and six subsequent rounds of propagation (P2-P7) that tested the effect of SEGS presence and vegetative propagation. Each experiment was conducted with three bioreplicates (independently inoculated plant lineages) per treatment. Best fit linear regression models included passage rounds, viral species (ACMV and EACMCV), and segment (DNA-A or DNA-B) nested in species (Tables 1 and 2, full experimental models are included in Tables S1 and S2.) Given the lack of evidence for a significant effect for several experimental treatments (replicate, temperature, and SEGS) and the failure of the full model to pass diagnostic tests of normality (Jarque-Bera test) and autocorrelation (Durbin-Watson test) (58), we grouped samples by generating a mean variant frequency. For example, to group SEGS treatments (no SEGS, + SEGS-1, + SEGS-2), we averaged allele frequencies across all three SEGS treatments for each SNP that shared the same passage, species, segment, and lineage. This had the effect of reducing the number of samples while generating a data set and model that passed diagnostic tests. Sample grouping consistent with this model was used for the rest of the analyses presented in this study. Both Veg2 and Veg6 grouped data were fit to the model:

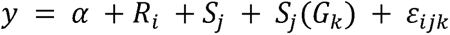

**Table 1.** Veg2, ANOVA results of nucleotide diversity as a response to experimental variables. Model R^2^ = 0.834. Model p = 0.00115.

|  | Degrees of Freedom | Sum of Squares | Mean Square | F | p |
| --- | --- | --- | --- | --- | --- |
| Species | 1 | 7.78 | 7.78 | 0.395 | 0.5496 |
| Species: Segment | 2 | 27.45 | 13.73 | 0.697 | 0.5296 |
| Passage | 1 | 658.75 | 658.75 | 33.453 | 0.0007 |
| Residual | 7 | 137.84 | 19.69 |  |  |

**Table 2.** Veg6, ANOVA results of nucleotide diversity as a response to experimental variables. Model R^2^ = 0.438. Model p = 0.00537

|  | Degrees of Freedom | Sum of Squares | Mean Square | F | p |
| --- | --- | --- | --- | --- | --- |
| Species | 1 | 231.4 | 231.4 | 2.06 | 0.165 |
| Species: Segment | 2 | 149.2 | 74.6 | 0.66 | 0.525 |
| Passage | 1 | 1632.6 | 1632.6 | 14.51 | 0.0009 |
| Residual | 23 | 2587.0 | 112.5 |  |  |

With *R* or the initial bombardment and each ith passage number, *S* species, *G* for segment nested in species, intercept *α* and error term *ε*. The overall models were significant for both Veg2 (*p* = 0.00115, *R*^2^ = 0.834) and Veg6 (*p* = 0.00537, *R*^2^ = 0.438). Only passage was significant in both Veg2 and Veg6, accounting for 94% and 79% of the variance explained by the model, respectively (as eta-squared calculated from Tables 1 and 2).

### Viral diversity changes over time

After determining that passage number (P1-P3 for Veg2; P1-P7 for Veg6) was the only factor that significantly impacted viral diversity quantitatively, we investigated how the diversity changed through successive rounds of vegetative propagation across the ACMV and EACMCV genome components. Sliding windows of π were calculated across segments accounting for their circular architecture and information retained after grouping read counts. π, which is the average pairwise difference between sequences in a sample (55), was examined on a population level and not by plant lineage, due to lineage failing to pass the test of normality and autocorrelation (Tables S1 and S2). Maximum π values were higher in the Veg6 experiment than the Veg2 experiment, suggesting that more rounds of vegetative propagation allowed for greater accumulation of nucleotide diversity (cf. Fig. 1 and 2). Comparison of P1, P2 and P3 between Veg2 and Veg6 revealed that the patterns of diversity are similar (Fig. 1a and c; S2a and c), except for a decrease in π in the ACMV *AV1* gene at P3 of Veg6 (cf. Fig. 1b and S2b). The patterns of increasing and decreasing π along genome components varied across passages, with several regions increasing in diversity across passages in Veg6 (Fig. S2a and S2c). Variation along the genome components was most apparent for EACMCV, which showed the highest level of nucleotide diversity in P7 for both genome components (Fig. 1c and S2c). The nucleotide diversity for EACMCV-A at P6 was intermediate between that of P7 and P4/P5, and the diversity of EACMCV-B was similar for P4-P6. In contrast, both genome components of ACMV displayed the highest levels of diversity at P4 and P6, with P5 and P7 showing lower levels (Fig. 2a and S2a).

**Fig 1.**
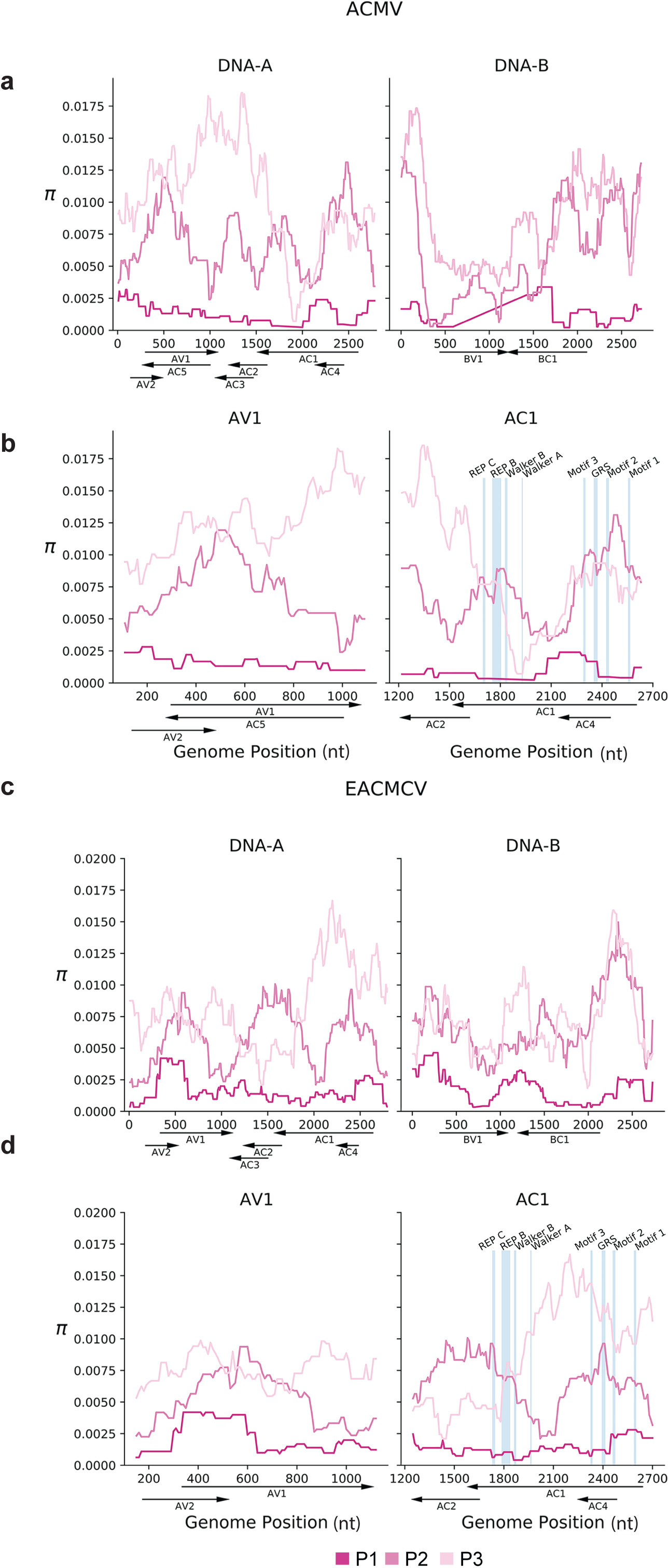
ACMV and EACMCV nucleotide diversity in the Veg2 experiment. (a) Sliding window analysis of nucleotide diversity (π) of ACMV DNA-A and DNA-B and (c) EACMCV DNA-A and DNA-B. Dark pink to lighter pink represents the nucleotide diversity across the genome of inoculated plants (P1) and two vegetative propagations (P2 and P3). Enhanced views of the nucleotide diversity of the *AV1* and *AC1* open reading frames during P1-P3 for ACMV-A (b) and EACMCV-A (d). Blue lines mark the locations of codons encoding functional motifs in the Rep protein, i.e. Rep C, Rep B, Walker B, Walker A (63), Motif 3 (62), GRS (64), Motif 2 (62) and Motif 1 (82). The motifs are shown to scale. Genome coordinates (nt), the positions of open reading frames and their directions of transcription are shown below each graph.

**Fig 2.**
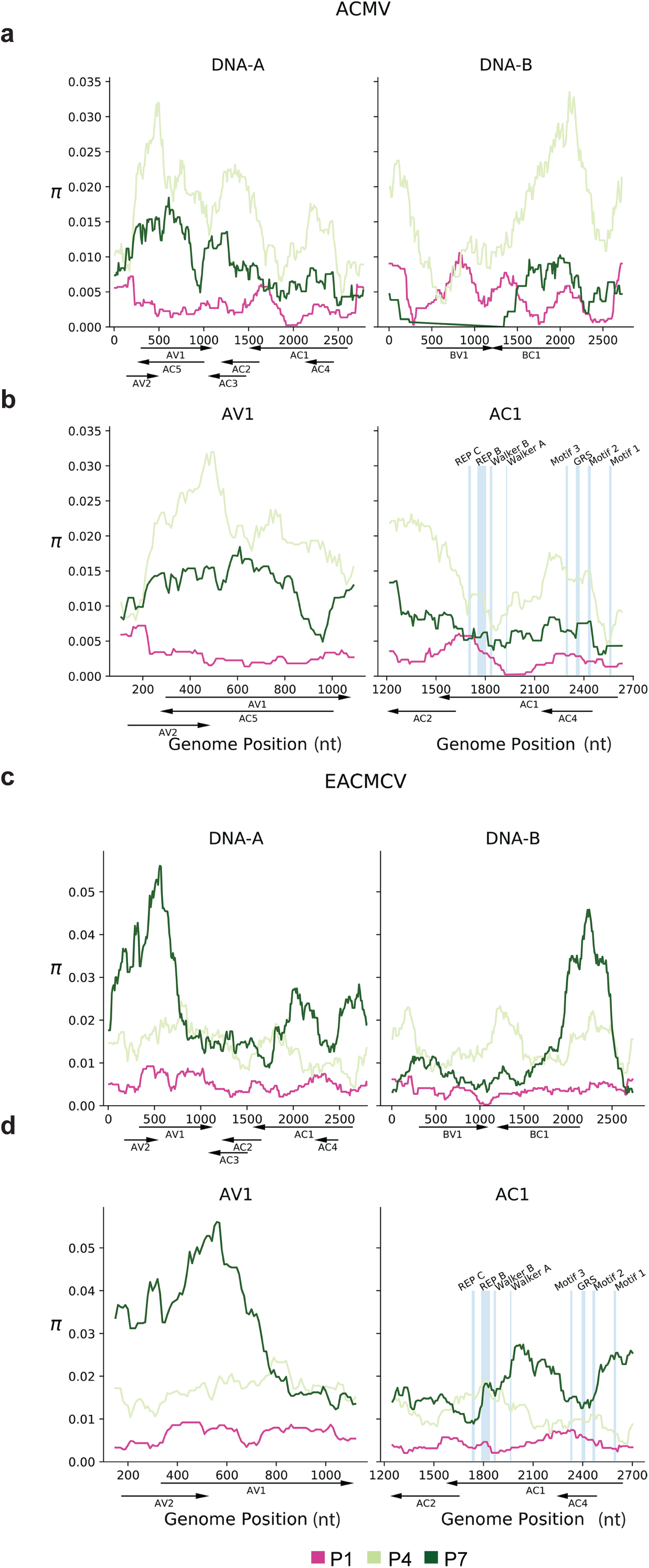
Veg6 ACMV and EACMCV nucleotide diversity sliding windows. (a) Sliding window analysis of nucleotide diversity (π) of ACMV DNA-A and DNA-B and (c) EACMCV DNA-A and DNA-B. Dark pink to dark green represents the nucleotide diversity across the genome of inoculated plants (P1) and two vegetative propagations (P4 and P6). Enhanced views of the nucleotide diversity of the *AV1* and *AC1* open reading frames during P1, P4, and P7 for ACMV-A (b) and EACMCV-A (d). Blue lines mark the locations of codons encoding functional motifs in the Rep protein, i.e. Rep C, Rep B, Walker B, Walker A (63), Motif 3 (62), GRS (64), Motif 2 (62) and Motif 1 (82). The motifs are shown to scale. Genome coordinates (nt), the positions of open reading frames and their directions of transcription are shown below each graph.

The DNA-A components of ACMV and EACMCV exhibited similar patterns of diversity in the passages showing the highest levels of nucleotide diversity, i.e., P4 for ACMV and P7 for EACMCV in the V6 experiment (Fig. 2a and c). The highest peaks of nucleotide diversity were over *AV2* and *AV1* (Fig. 2a and c). In ACMV-A, nucleotide diversity covered a broader region of *AV1* overlapping with *AC5* encoded on the opposite strand. Peaks of diversity were also observed over other overlapping regions of ACMV-A: *AC1*/*AC4* and *AC2*/*AC3* (Fig. 2a and 2b). In EACMCV, two peaks of diversity occurred over non-overlapping coding regions of *AC1,* with overlapping regions showing moderate diversity (Fig. 2c and 2d). It was unexpected that nucleotide diversity appeared to peak in regions where viral genes overlap in different reading frames, in which codon wobble positions would be constrained. Chi-square tests confirmed that diversity was not constrained by the number of overlapping protein coding regions within ACMV-A or EACMCV-A (p > 0.21, Table S3-3). The DNA-B components of ACMV and EACMCV also displayed similar patterns of nucleotide diversity (Fig. 2a and 2c). High levels of diversity were seen in the 5’ intergenic region, and *BC1* had higher levels compared to *BV1*.

### Test of evolution/demographic changes

After examining viral diversity on the genomic level, we examined the potential impact of more frequent individual variants. We examined the effect of varying minimum variant frequency thresholds on Tajima’s D, which infers selection and/or demographic events (population size changes not due to selection) and is sensitive to rare variants (59). Tajima’s D is the standardized difference of two different ways to calculate the expected nucleotide diversity. The caveat with using Tajima’s D is that purely demographic changes in the population can result in extreme values of Tajima’s D. Our study has an advantage to this point over the problem of disentangling selection and demographic histories in wild populations: given our experimental design we have a good understanding of what demographic effects are possible. We also do not use dN/dS as this measure is not well suited to the short evolutionary time frame and likely small population sizes found in these experiments. We filtered SNPs by varying the minimum frequency for each of the Veg2 and Veg6 experiments (Fig. 3). In Fig. 3, SNP frequencies were grouped by passage and segment nested within species (ACMV-A, ACMV-B, EACMCV-A, EACMCV-B) for each experiment. General interpretations of Tajima’s D typically hold for minimum variant frequencies of 2-4% for SNP analysis (59). Hence, we used a minimum variant frequency of 3% in our analysis (Fig. 3, dotted vertical line) to rule out the influence of very rare variants and simplify the analysis of potential functional implications of variance. In the Veg2 experiment, Tajima’s D > 2 was significant (P<0.01) in P2 for ACMV DNA-B and in P3 for ACMV DNA-A, ACMV DNA-B, and EACMCV DNA-B (Fig. 3a and b), but not for EACMCV DNA-A in any passage (Fig. 3b). In contrast, the later passages of the Veg6 experiment were positively significant for all segments (Fig. 3c and d), but the magnitudes of Tajima’s D for all segments varied across passages in a nonlinear fashion similar to the nucleotide diversity profiles (Fig. S2). The marginal significance (0.01 < *P* < 0.05) of a positive Tajima’s D in the Veg2 and Veg6 experiments indicates more intermediate-frequency variants than expected and does not support strong selective sweeps in either experiment. We note our data do not support strong selective sweeps or detectable population bottlenecks even though our methods are not as conservative as those in other pool-seq genetic diversity analytics (56).

**Fig 3.**
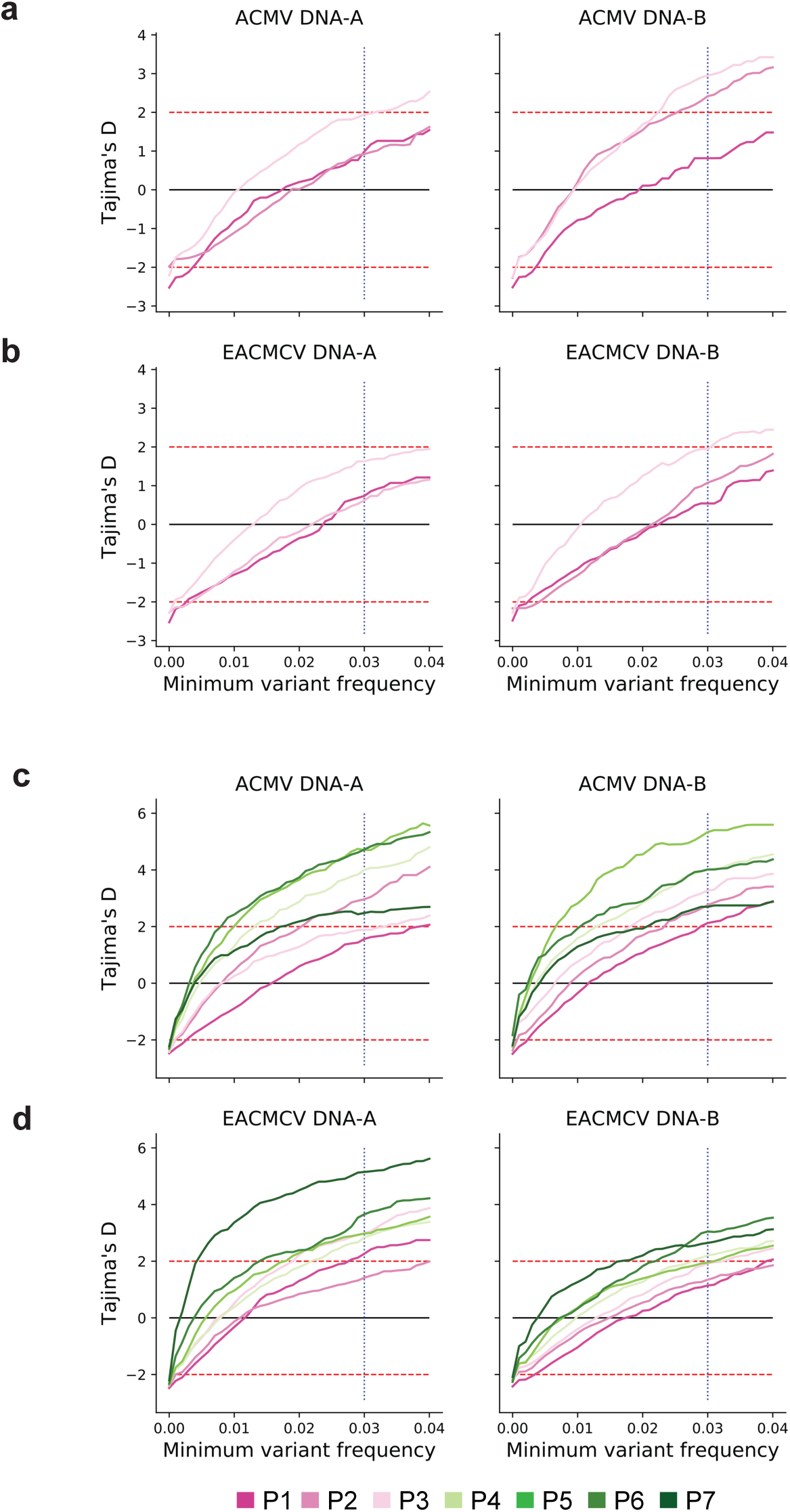
Tajima’s D analysis by passage and minimum variant frequency cutoff. Tajima’s D analysis passage (P) for (a) ACMV DNA-A and DNA-B and (b) EACMCV DNA-A and DNA-B in the Veg2 experiment. Tajima’s D by passage for (c) ACMV DNA-A and DNA-B and (d) EACMCV DNA-A and DNA-B in the Veg6 experiment. Dotted red horizontal lines represent thresholds for significant Tajima’s D values (59). The vertical blue dotted line represents the minimum variant frequency of 3% used in this study. The lines representing each passage are color coded, as shown at the bottom.

### Biases in nucleotide substitutions

High mutation rates of ssDNA viruses have been attributed to deamination and oxidative damage (10, 60). Thus, we investigated whether nucleotide substitution bias was detectable within the timeframe of our experiments. We accounted for multiple testing by the Benjamini-Hochberg method with FDR=0.01 (61). Substitutions were biased toward C→T and G→A for all components in both experiments (p < 10^−6^). A bias toward C→A was observed in both experiments and toward G→T for all but ACMV DNA-B in the Veg6 experiment (p < 10^−3^, see Table S3-4).

### Changes in viral diversity at the codon level

To examine the impact of nucleotide diversity on amino acid codons, we looked at SNPs that passed our 3% threshold, occurred in both technical replicates, and were present in more than one passage (see Table S3-5 (Veg2) and S3-6 (Veg6) for a list of all of the SNPs found in at least two passages). ACMV had 125 codon changes, while EACMCV had 97 changes, with codon changes occurring in all ORFs of both viruses (Fig. 4a and b). When adjusted for ORF length, *AC1* of both viruses and ACMV *BV1* contained the fewest SNPs, with the other ORFs showing similar levels of SNPs (Fig. 4c and d). We also observed more non-synonymous SNPs than synonymous across both genomes (ACMV: N:80 S:49; EACMCV N:54 S:42). SIFT analysis predicted that the proportion of amino acid substitutions that negatively impact protein function was greater for EACMCV (65%) than for ACMV (52%, Table S3-6). Detrimental amino acid substitutions were predicted to occur in all viral ORFs. Some ORFs had higher number of non-synonymous mutations, with *AV2*, *AC4*, *AC5* of ACMV and *AC4* of EACMCV having the highest (Fig. 4a and b). When adjusted for ORF length, *AC1* and *BV1* had the lowest numbers and fraction of non-synonymous changes for both viruses. Rep is a complex enzyme that catalyzes multiple reactions necessary for viral replication (28) and, as such, may be less tolerant to amino acid substitution. In addition to viral DNA trafficking, NSP is engaged in a number of host interactions that may be sensitive to amino acid changes (40).

**Fig 4.**
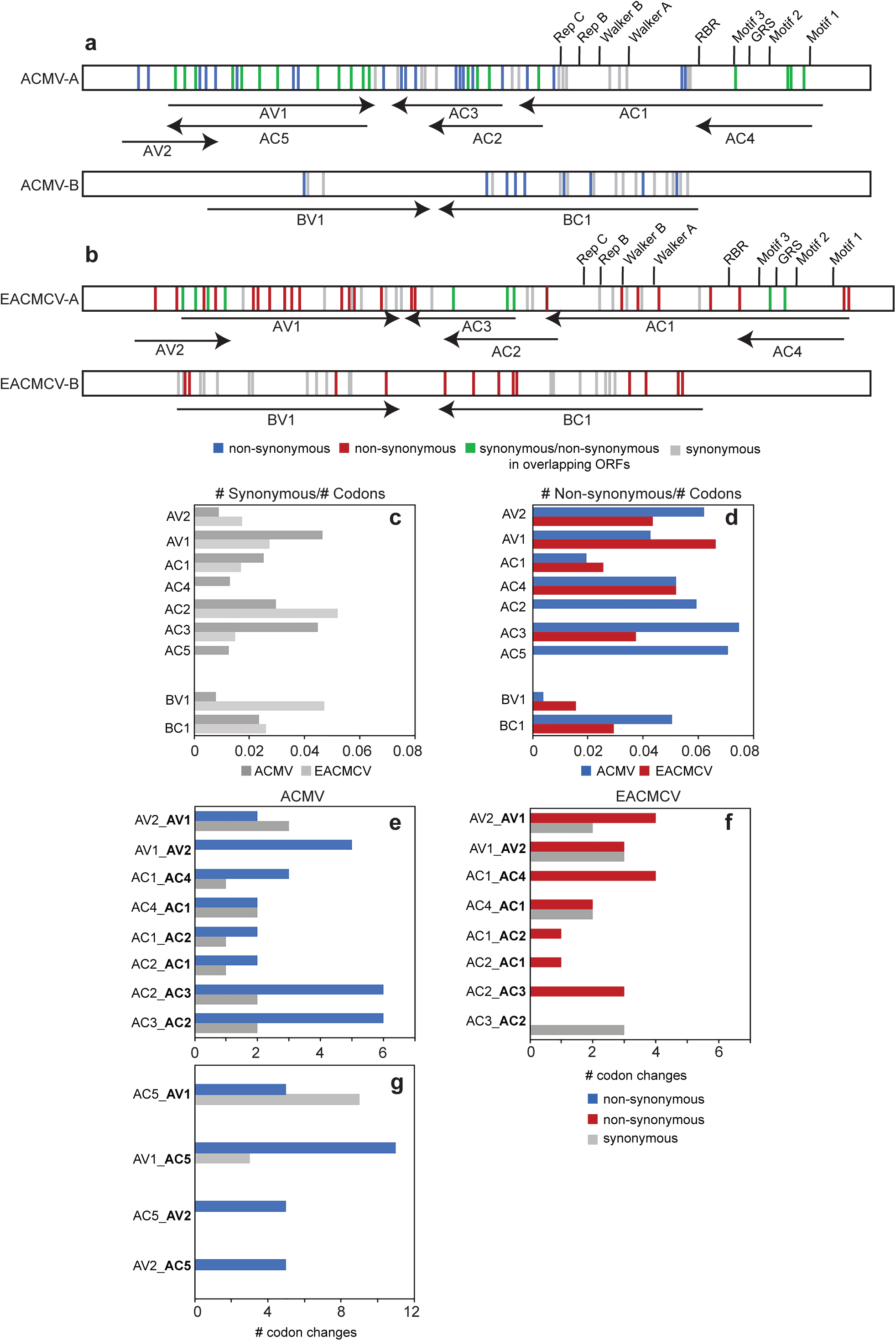
Synonymous and non-synonymous codon changes in ACMV and EACMCV. Diagram of the locations of synonymous codon changes (grey), unassigned changes in overlapping open reading frames (ORFs; green), and non-synonymous changes for (a) ACMV (blue) and (b) EACMCV (red). The number of synonymous (c) and non-synonymous (d) codon changes normalized to the total number of codons in the ORF for ACMV (dark grey and blue) and EACMCV (light grey and red). The number if synonymous (grey) and non-synonymous (ACMV-blue, EACMCV-red) codon changes in overlapping coding sequences in ORFs overlapping within ACMV excluding *AC5* (e), EACMCV (f), and regions within ACMV that overlap *AC5* (g). The overlapping ORF is designated first and the ORF assessed for the codon change is designated second in bold.

We asked how the SNPs impacted codons in known functional domains and motifs of the Rep protein (annotated in Fig. 1 and 2). The *AC1* ORFs of both viruses include long stretches devoid of polymorphisms, and the majority of changes were synonymous. ACMV *AC1* had synonymous SNPs in the DNA cleavage Motif 3 (62) and the Walker A helicase motif, while EACMCV *AC1* had a synonymous change in the Walker B helicase motif (63). The sequence of the EACMCV *AC2* promoter element that overlaps the Walker B motif was maintained in the mutant Tyr257Tyr (63). However, there were several non-synonymous changes distributed throughout the DNA binding/cleavage, oligomerization, and DNA helicase domains of their Rep proteins (29) (Table S3-6). Most notably, there was a SNP in EACMCV *AC1* that resulted in a Phe75Val codon change at a highly conserved position in the GRS motif (64).

We also looked at the potential effects of codon changes in other viral proteins. More SNPs were associated with ACMV *AC2* than EACMCV *AC2* (ACMV: 15, EACMCV: 5, Fig. 4). The amino acid changes were distributed across regions of the *AC2* protein associated with DNA binding (65), suppression of PTGS (66), and transcriptional activation (67). ACMV *AC3* was also associated with more SNPs the EACMCV *AC3* (ACMV: 16, EACMCV: 7, Fig. 4), and most of the SNPs in ACMV *AC3* resulted in non-synonymous codons changes, consistent with the capacity of REn to accommodate amino acid substitutions at many positions throughout the protein (68). The *AC3* gene contained the only codon change with a SNP in both viral genomes, i.e., Pro77 changing to Tyr in ACMV and to His in EACMCV. In contrast, the number of SNPs in *AV1* was similar for both viruses, with approximately half of the SNPs resulting in codon changes. The non-synonymous codon changes impacted amino acids associated with CP nuclear localization, multimerization, and ssDNA binding (69), as well as whitefly transmission of begomoviruses (70–72). There were non-synonymous changes throughout the *BC1* gene of both viruses, some of which introduced amino acid changes at positions in the MP implicated in oligomerization (73) and subcellular targeting (74). One of the few non-synonymous codon changes in *BV1* was in the NSP nuclear export signal (75).

We also examined codon changes in regions where ORFs overlap on DNA-A. ACMV had more non-synonymous codon changes than synonymous ones in the region of *AV1* that overlaps with *AV2* and *AC5* (Fig. 4e and g), but no such bias was observed in the *AV2*/*AV1* overlapping region in EACMCV (Fig. 4f). Both viruses showed a tendency for non-synonymous changes to accumulate in the *AC4* ORF (Fig. 4e and f), which occurs entirely within *AC1*. Similar results have been reported for Tomato leaf deformation virus and Tomato yellow leaf curl virus (76, 77). ACMV *AC2* and *AC3* both had a higher proportion of non-synonymous codon changes in their overlapping region (Fig. 4e), but this was not seen for EACMCV, which had non-synonymous codon changes in *AC3* but not *AC2* (Fig. 4f).

Some of the SNPs detected in the Veg6 experiment are also present in a historical database of CMB sequences available from GenBank (Table S3-7), as ascertained by querying recently described multiple alignments (Crespo et al., in preparation). Collectively, 19 Veg6 SNPs for ACMV DNA-A occurred 451 times in the set of 851 DNA-A sequences in GenBank (http://zenodo.org/record/4029589). Six EACMCV DNA-A Veg6 SNPs occurred 20 times in the same set of sequences, 3 ACMV DNA-B SNPs occurred 5 times (in a set of 104 sequences; http://zenodo.org/record/3964979), and 4 EACMCV DNA-B SNPs occurred one time each (in a set of 243 sequences; http://zenodo.org/record/3965023) (78, 79). The SNPs represented in the historical database represent 9% of the total SNPs identified in the Veg6 study with 15%, 6%, 8%, and 5% of the total SNPs identified for ACMV-A, ACMV-B, EACMCV-A, and EACMCV-B, respectively. This indicates that the majority of the SNPs observed in our studies are novel. The SNPs observed in our experiments, those currently in the historical database occurred at a higher allele frequency than those not represented in the historical database for ACMV-A at P 3-7, ACMV-B at P5-6, EACMV-A at P5 and P7, and EACMCV-B at P5 (P-value <0.05, Table S3-8). While we cannot rule out random genetic drift, these results beg further investigation of SNPs found in nature where, at certain passages may have been under positive selection in our experiments (Table S3-8). Yet, a majority of the changes found in the historical database for each genome segment were non-synonymous, except for in EACMCV-A where roughly equal numbers of synonymous and non-synonymous changes were observed (Table 3). Of the non-synonymous changes, 40% were predicted to be detrimental based on SIFT analysis (53) (Table S3-7).

**Table 3.** Number of SNPs observed in the historical database for Veg6 experiment by genome component.

|  | ACMV-A | ACMV-B | EACMCV-A | EACMCV-B |
| --- | --- | --- | --- | --- |
| Synonymous | 5 | 1 | 3 | 0 |
| Non-Synonymous | 14 | 2 | 3 | 4 |

### Changes in common region sequences

We examined nucleotide changes that occurred during multiple passages in the noncoding sequences of DNA-A and DNA-B. There were 17 and 19 changes in the 5’ intergenic sequences of ACMV DNA-A and DNA-B, respectively, while EACMCV had 12 and 35 changes in DNA-A and DNA-B, respectively (Table S3-6). We also observed 7 nucleotide changes in the 3’ intergenic region of EACMCV-B, but none for the other viral components, which all have very short 3’ intergenic regions with bidirectional polyadenylation signals (80).

The 5’ intergenic regions of DNA-A and DNA-B contain a conserved sequence designated as the common region that contains the origin of replication and divergent promoter sequences (Fig. 5 and 6). Changes in the common region occurred primarily outside of known conserved cis elements involved in replication (iterons and the stem loop (26, 27)) and transcription (TATA box, CLE element and CCAAT box (27, 81)). However, we observed an A→G change in the putative TATA box of the ACMV *AV2* promoter (Table S3-6). We also detected G→C transversions in the stem sequences downstream of the nick site of both ACMV DNA-A and ACMV DNA-B (Fig. 5b), but we did not find compensatory changes in the upstream stem sequences. Given that the stem structure is necessary for viral replication (26), it was surprising that the 3’-stem SNPs were maintained in the population at a frequency of ca. 5% in at least 2 passages of the Veg6 experiment. We observed a G→A transition at position 2626 in EACMCV-B. Comparison to the historical database indicated that the 5 G residues starting at position 2626 constitute a third conserved iteron in the origin of replication (Fig. 6b; Table S3-7). However, the G at position 2626 is followed by 5 G residues, suggesting they can also serve as an origin iteron when G-2626 is mutated (82).

**Fig 5.**
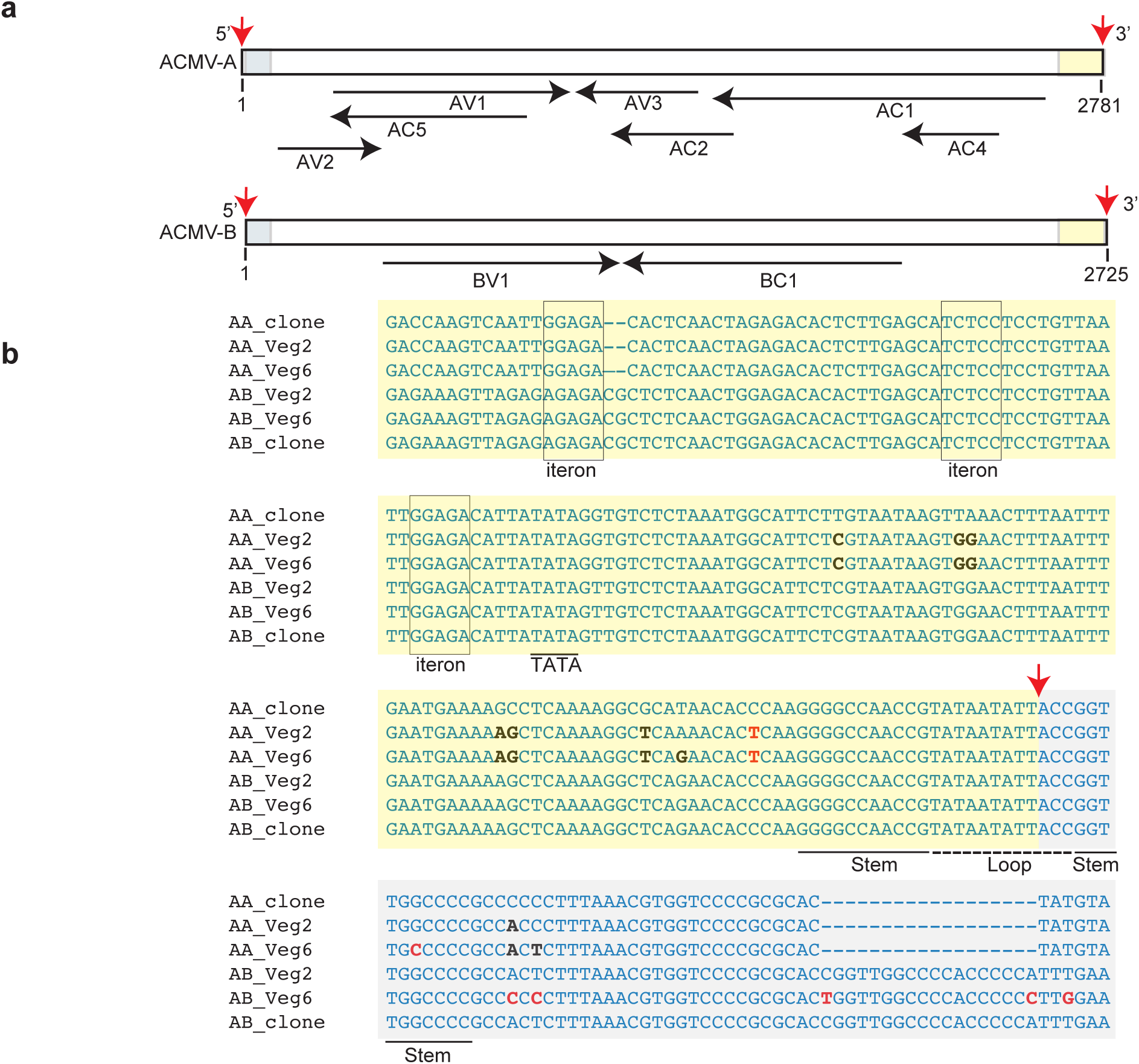
ACMV common regions undergo sequence convergence. (a) Linear maps of ACMV DNA-A and ACMV DNA-B. The maps were linearized in the common region at the cleavage site in the top strand of the viral origin of replication. Red arrows mark the 3’-OH and the 5’-P of the nick site (99). The common region upstream (yellow) and downstream (grey) of the nick site are marked. The open reading frames and directions of transcription are shown by the black arrows below. (b) ACMV DNA-A and ACMV DNA-B sequences showing their common regions in the circularized genomic form. The labeling is the same as in (a), with the nick site indicated by a red arrow and the upstream and downstream sequences marked by yellow and grey shading, respectively. The iterons (boxed) and the hairpin motif (stem: underlined; loop: dotted line) involved for the initiation of viral replication are marked. The TATA box (underlined) for complementary sense transcription is also labeled. SNPs showing convergence of the common region sequences of DNA-A and DNA-B are in black typeface, and other SNPs are in red typeface.

**Fig 6.**
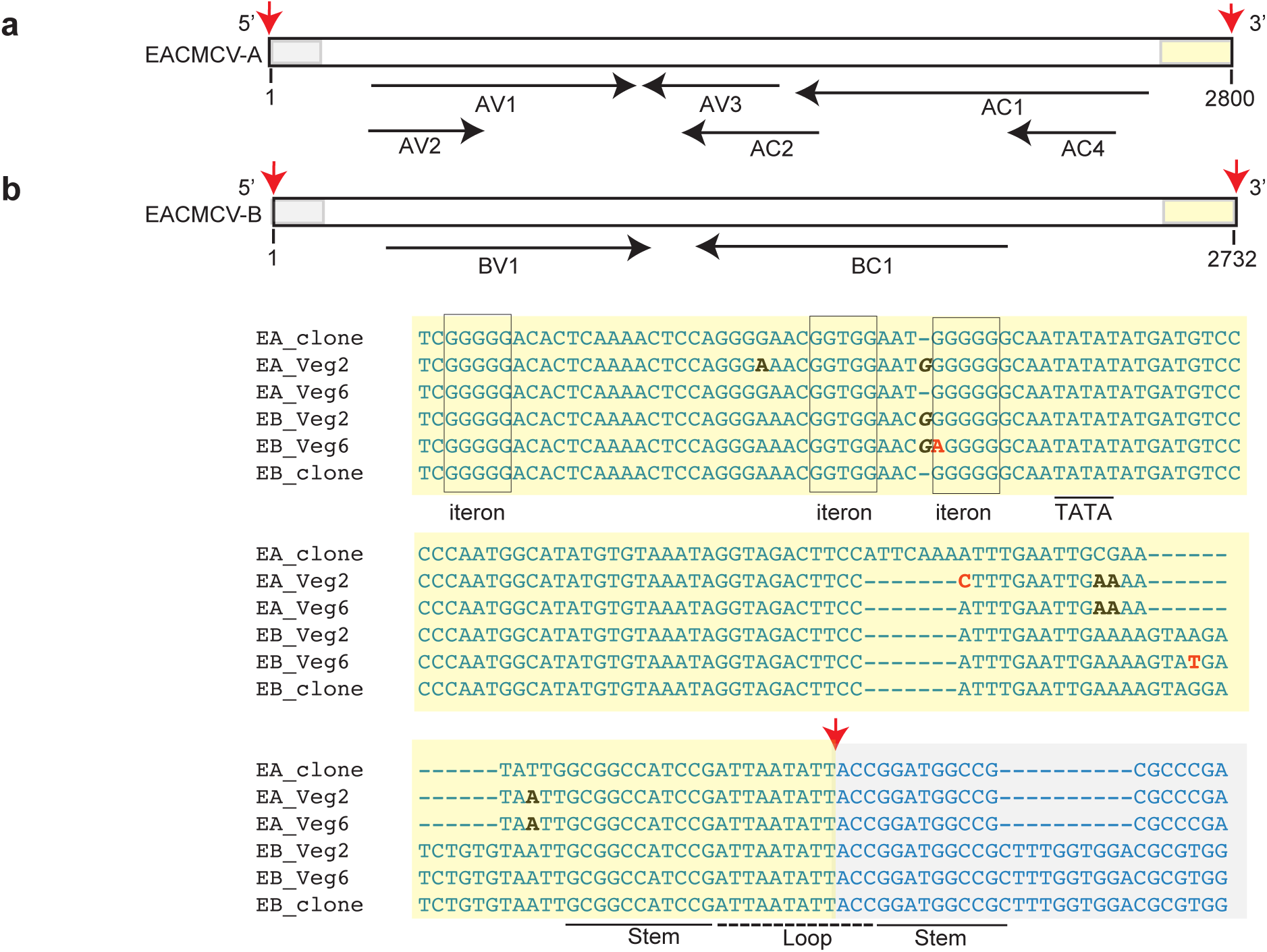
EACMCV common regions undergo sequence convergence. (a) Linear maps of EACMCV DNA-A and EACMCV DNA-B. The maps were linearized in the common region at the cleavage site in the top strand of the viral origin of replication. Red arrows mark the 3’-OH and the 5’-P of the nick site (99). The common region upstream (yellow) and downstream (grey) of the nick site are marked. The open reading frames and directions of transcription are shown by the black arrows below. (b) EACMCV DNA-A and EACMCV DNA-B sequences showing their common regions in the circularized genomic form. The labeling is the same as in (a), with the nick site indicated by a red arrow and the upstream and downstream sequences marked by yellow and grey shading, respectively. The iterons (boxed) and the hairpin motif (stem: underlined; loop: dotted line) involved for the initiation of viral replication are marked. The TATA box (underlined) for complementary sense transcription is also labeled. SNPs showing convergence of the common region sequences of DNA-A and DNA-B are in black typeface, and other SNPs are in red typeface. Insertion is indicated by italics.

We also found that the common regions of the DNA-A components of both ACMV and EACMCV acquired mutations such that they more closely resembled the sequences of their cognate DNA-B common regions (Fig. 5b and 6b). This occurred in all bioreplicates of each passage in both Veg2 and Veg6 experiments. The ACMV DNA-A common region had SNPs at 11 positions of which 9 matched ACMV-B (Fig. 5b). ACMV DNA-B had 5 SNPs, 3 at equivalent positions as ACMV-A substitutions. The EACMCV DNA-A common region had 5 SNPs with 4 matching EACMCV DNA-B, and EACMCV DNA-B had 2 other SNPs (Fig. 6b).

We observed multiple deletions in the alignments of the ACMV DNA-A and EACMCV DNA-A common regions, while their cognate B components maintained the same length without indel mutations throughout all of the propagation experiments. There was a 7-bp deletion in EACMCV DNA-A in both the Veg2 and Veg6 experiments that resulted in a sequence match with EACMCV DNA-B, another mutation that made the DNA-A intergenic region better resemble that of the cognate DNA-B (Fig. 6b). The deletion was accompanied by CG→AA changes 11-12 bp downstream. Mapping of the paired-end reads established that the 7-bp deletion and associated nucleotide changes in EACMCV DNA-A were not due to erroneous mapping of EACMCV DNA-B reads. Moreover, the deletion and nucleotide changes were not detected by Illumina sequencing of the plasmid controls, confirming that the variant was not present at low frequency in the EACMCV DNA-A inoculum. Together, these results suggested that the intergenic regions of ACMV DNA-A and EACMCV DNA-A sequences were less fit than those of their DNA-Bs, and there was selective pressure causing convergent evolution through inter-segment recombination with the more optimal sequences that resemble their cognate DNA-B sequences.

## Discussion

Viruses exist as populations of related sequence variants (15). This reservoir of genetic diversity enables plant viral populations to change rapidly in response to environmental conditions, agricultural practices, and different hosts (17–20). Thus, understanding viral diversity and the external parameters that impact diversity is essential to develop durable disease resistance strategies. We characterized the genetic diversity of ACMV and EACMCV after inoculation of cassava plants with cloned viral sequences. We examined the effects of vegetative propagation as an agriculture practice, temperature, and the presence of exogenous DNA sequences on the nucleotide diversity of the two CMBs. Our studies revealed that repeated vegetative propagation of infected cassava increased genome-wide nucleotide diversity of CMBs without detectable bottlenecks. Our experimental design starting from defined clones allowed us to confidently perform fine-scale analyses of genetic diversity using π, and to detect signatures of selection using Tajima’s D involving large population sizes.

In our primary analysis of environmental factors, we found that vegetative propagation had the largest, and only significant, impact on CMB nucleotide diversity (Table 1 and 2). Multiple rounds of vegetative propagation of potato tubers infected with potato virus Y (PVY) also resulted in an overall increase in nucleotide diversity, and vegetative propagation had a larger effect on PVY diversity than vector transmission (83). Diversity profiles through passages varied depending on the PVY strain, with some strains increasing linearly and other strains peaking after the first vegetative passage and then decreasing (83). Our studies uncovered differences between CMB species, in that ACMV nucleotide diversity peaked at P4 and P6 and then decreased, while EACMCV diversity increased linearly through passages (Fig. 2 and Fig. S2a). Vegetative propagation was also the main factor leading to recombination events and generation of new geminivirus species in sweet potato (84, 85). Together, these results show how transmission mode impacts viral populations and evolution in root and tuber crops. Our results underscore the importance of discarding infected plants and providing access to virus-free planting material instead of using vegetative propagation to reduce virus spread and the emergence of new viral variants.

Surprisingly, increasing temperature had no significant impact on CMB nucleotide diversity (Supplementary Table 2). A 10°C shift has been associated with an increase in potexvirus nucleotide diversity in tomato, and the selection of SNPs in a strain-dependent manner (86). We used a 2°C temperature shift, based on predicted increases for global warming in Africa by 2030 (47). The different outcomes in the two studies are likely due to the 5-fold difference in the temperature shifts but could also reflect differences in how DNA vs. RNA viruses or cassava vs. tomato plants respond to elevated temperatures. Nevertheless, our results suggest that the predicted 2°C temperature shift in Africa, may not be a main driver of CMB diversity (47).

Some begomovirus satellites have been associated with elevated levels of viral DNA and suppression of host DNA methylation and silencing pathways (87), and as such have the potential to impact viral diversity. Although it is not known if SEGS-1 or SEGS-2 function in a similar manner, they have several features in common with satellites (46). We were unable to detect an effect of either SEGS-1 or SEGS-2 on viral nucleotide diversity in coinoculation experiments with CMBs (Table S3-1). However, these results do not rule out that other types of begomovirus satellites may impact viral diversity.

We used Tajima’s D to examine the patterns of nucleotide diversity through time and to gain insight into whether the CMB genomes were undergoing selection during vegetative propagation. Tajima’s D compares the average number of pairwise differences with the number of segregating sites and determines selection or bottleneck pressure based on deviations from constant population size. Our analysis uncovered a significant positive change in variant frequency for both ACMV and EACMCV (Fig. 3), indicative of an increase in the number of variants during vegetative propagation. However, the marginal significance is consistent with the generation of a population of intermediate variants from a founder event, i.e., inoculation of a cloned viral sequence, and that most variants are not under positive selection. The lack of selection pressure is consistent with the high level of variation in the amount of nucleotide diversity across passages (Fig. 2 and Fig. S2a). However, the intermediate variant population may undergo selection with more time, more passages, or most importantly, if the diverse population is exposed to a novel selection pressure. A similar phenomenon was observed for PVY SNP populations associated with 5 rounds of vegetative propagation of field-grown, infected potato (da Silva et al., 2020). These studies found that very few variants became fixed in the PVY population due to positive or negative selection, indicating that a small number of viral variants contributed to each new population after propagation (83). In our study, the variation in nucleotide diversity patterns across the CMB genomic components from one passage to the next (Fig. S2a) is consistent with a small fraction of viral variants propagated to the next generation with no mutations fixed in the population for more than 3 passages.

We observed more non-synonymous changes than synonymous changes in the genomes of both ACMV and EACMCV (Fig. 4c and d), which is consistent with the higher possibility of non-synonymous mutations in coding regions by random chance. However, a large fraction of the non-synonymous codon changes also observed in both viruses were predicted to cause detrimental amino acid substitutions using SIFT analysis, which has been correlated with a lack of purifying selection (88). The strongest effects of purifying selection (fewest non-synonymous changes, large stretches without changes) was observed in AC1, not *AV1* as has been observed in field isolates of EACMV (10). This difference is likely a direct result of the vegetative transmission mode being tested here; many plant viruses have strong purifying selection on their capsid proteins because of their interactions with both plant and insects (89). During vegetative propagation of CMVs there is no selective pressure from a whitefly vector, and we see that relaxation in the higher numbers of non-synonymous changes in *AV1* compared to *AC1*. While these results rule out strong purifying selection acting on CMB viral ORFs, we cannot rule out balancing selection within the timeframe of our studies based on the results of Tajima’s D. The presence of SNPs from our experiment in field-collected sequences in GenBank means that some of the variation we observe is not so deleterious as to never be isolated in nature, but we saw no evidence that this subset of SNPs was under more positive selection in our vegetative passaging. Because field populations of CMBs are not exhaustively sequenced, it remains unclear what fraction of the SNPs in this experiment have and would persist and thrive in the field.

Unlike the viral ORFs, we observed multiple nucleotide substitutions in the common regions of the A components of ACMV and EACMCV that were present in all bioreplicates, resulting in sequences that more closely resembled the common regions of their cognate B components. The common region includes cis-acting sequences that are necessary for replication. When an infectious clone is made, a single sequence is cloned out of the viral population in the source plant, and that sequence must have a functional origin to be viable infectious clone. This is in stark contrast to viral sequences carrying detrimental codon mutations, which can be complemented in trans by other viral components carrying functional promoters and ORFs (90). However, the cloned sequence may not have an efficient origin and, thus, would be under selective pressure to better compete with more efficient origin sequences. This type of competition was seen in origin mutants of Tomato golden mosaic virus (TGMV) DNA-B in protoplast replication assays (26). Hence, the convergence of CMB DNA-A common region sequences toward DNA-B may reflect evolutionary selection of a more efficient origin through sequentially acquired intersegment recombination, which would explain the multiple similar substitutions more easily than sequential acquisition of independent mutations. Examples of intersegment (inter-component) recombination of common regions have been observed during passaging in *Nicotiana benthamiana* plants under laboratory conditions with tomato mottle virus, bean dwarf mosaic virus, and African cassava mosaic virus (91, 92), but the directionality has previously always been a DNA-B common region becoming more like DNA-A. Intersegment recombination of the common region has also been frequently observed in the ssDNA phytopathogenic nanoviruses, which have six or more genome components (93).

Nucleotide substitution bias has been observed for geminivirus sequences in historical datasets (10) and in experimental populations examined within single plants infected for up to five years (94, 95). Our results showed that nucleotide substitution biases are readily detected in a 2-month timeframe for ACMV and EACMCV, and that this short time is sufficient to detect nucleotide substitution biases in both transitions (C→T and G→A), consistent with historical sequences of East African cassava mosaic virus (10) and experimental infection of Tomato yellow leaf curl China virus (95) and one transversion (G→T, previously reported for experimental populations (94)) for both viruses. It has been proposed that the nucleotide substitution bias is due to oxidative damage (10, 94). The repeated pattern of nucleotide substitution bias in our experiments points to oxidative damage (96). The C→T and G→A transitions could be due to hydroxylation mediated by reactive oxygen species, possibly by oxidative deamination of cytosine in the case of C→T mutations (97), and the G→T transversions could be from oxidative damage converting guanine to 8-oxoguanine, which base pairs with adenine leading to a G-C base pair mutating to a T-A base pair (98). Our SNP nucleotide biases match the substitution biases of ssDNA virus historical data sets, showing that the same molecular patterns can be observed over months as decades.

This study provides evidence that farming practices, such as vegetative propagation, can have a large impact on genetic diversity across CMB genome components. This study also presents evidence that CMB genes are under relaxed selection pressure during vegetative propagation, with *AC1* and *BV1* under the strongest purifying selection pressure. This is not the case for the intergenic region, especially in the convergence of the DNA-A common region sequences toward DNA-B, likely reflecting selection of a more efficient origin. This pattern is consistent with mutated viral genomes being able to complement protein functions in trans, but there being strong selection for optimal origin of replication function, which cannot be complemented. Understanding how CMBs and other ssDNA viruses are evolving through experimental evolution studies such as this will ultimately help inform and improve strategies for disease management.

## Abbreviations

ACMV: African cassava mosaic virus
EACMCV: East African cassava mosaic Cameroon virus
CMD: Cassava mosaic disease
CMB: Cassava mosaic begomovirus
PTGS: Post-transcriptional gene silencing
TGS: Transcriptional gene silencing
ssDNA: Single stranded DNA
dsDNA: Double-stranded DNA
RCA: Rolling circle amplification
CP: Coat protein
Rep: Replication-associated protein
REn: Replication enhancer protein
MP: Movement protein
NSP: Nuclear shuttle protein
SEGS: Sequence enhancing geminivirus symptoms

## Acknowledgments

We thank Mary Beth Dallas for her help growing cassava plants. Infectious clones were generously provided by Vincent Fondong (Addgene plasmids 159134 to 159137; https://www.addgene.org/browse/article/28211870/). We thank the NC State University Genome Science Laboratory for their support. The Cassava Virus Evolution Galaxy server (https://cassavavirusevolution.vcl.ncsu.edu/) is hosted in the NC State University Virtual Computing Lab, and we thank Matthew Gronke, Saahil Chawande, and Jim White for server setup and maintenance. This work also used the Amarel cluster maintained by the Office of Advanced Research Computing (OARC) at Rutgers, The State University of New Jersey.

## Author contributions

C.A. and E.L. were responsible for writing and preparing the manuscript, experimental design, collection, and data analysis. J.S.H. was responsible for experimental design and data analysis. D.D. and L.M.-Y. conducted library preparation. L.H.-B., S.D., A.L.J, I.C., and G.G.K. were responsible for experimental design and manuscript preparation.

## SUPPLEMENTAL MATERIAL

**S1. ACMV and EACMCV clones.** Diagram of A and B components of ACMV (a, b) and EACMCV (c, d). Black arrows represent opening reading frames, and the red arrow indicates nick site. (e) The table shows the gene name of each opening reading frame, the corresponding protein name, and function(s) of the protein in infection.

**S2. Veg6 ACMV nucleotide diversity sliding windows of all 7 rounds.** (a) Sliding window analysis of nucleotide diversity (π) of ACMV DNA-A and DNA-B and (c) EACMCV DNA-A and DNA-B. Dark pink to dark green represents the nucleotide diversity across the genome of inoculated plants (P1) and six vegetative propagations (P2-P7). Enhanced views of the nucleotide diversity of the *AV1* and *AC1* open reading frames during P1-P7 for ACMV-A (b) and EACMCV-A (d). Blue lines mark the locations of codons encoding functional motifs in the Rep protein, i.e. Rep C, Rep B, Walker B, Walker A (63), Motif 3 (62), GRS (64), Motif 2 (62) and Motif 1 (82). The motifs are shown to scale. Genome coordinates (nt), the positions of open reading frames and their directions of transcription are shown below each graph.

**S3. Supplementary Tables 1-7**

Table S3-1. Veg6 full experimental model ANOVA results.

Table S3-2. Full experimental model ANOVA results for Veg2 experiment.

Table S3-3. χ2 analysis of the number of coding regions at a genome position and codon changes.

Table S3-4. Analysis of nucleotide substitution bias in the Veg2 and Veg6 experiments.

Table S3-5. Nucleotide changes at a frequency >3% in multiple passages in the Veg2 experiment.

Table S3-6. Nucleotide changes at a frequency >3% in multiple passages in the Veg6 experiment.

Table S3-7. Veg6 nucleotide changes present in the historical database.

Table S3-8. Mann-Whitney test results between frequency of SNPs in Veg6 and those found in the historical database.

