## Supplemental Figure 1 for "Population diversity of cassava mosaic begomoviruses increases over the course of serial vegetative propagation"

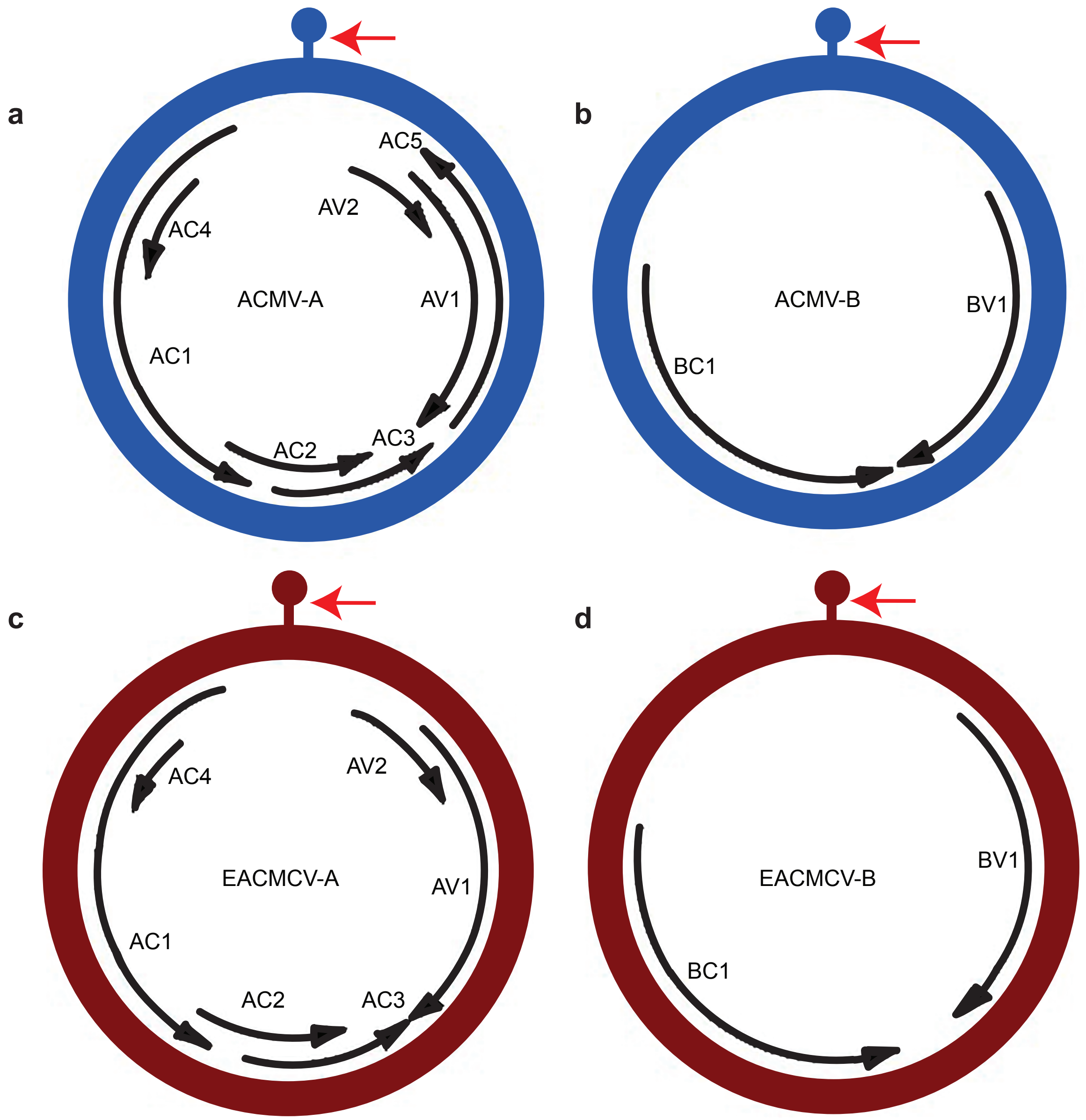


e

| Gene Name | Protein Name | Major Function |
|---|---|---|
| AV2 | AV2 | Inhibitor of post-transcriptional gene silencing (PTGS) |
| AV1 | CP | Encapsidation, whitefly transmission |
| AC5 | AC5 | Unknown function |
| AC3 | REn | Viral replication |
| AC2 | TrAP | Viral transcription, viral supressor of silencing (VSR) |
| AC1 | Rep | Viral replication |
| AC4 | AC4 | VSR |
| BC1 | MP | Movement protein |
| BV1 | NSP | Nuclear shuttle protein, anti-defense protein |
