## Supplementary figures and images for "Population diversity of cassava mosaic begomoviruses increases over the course of serial vegetative propagation"

### Supplemental Figure 2

S2 Aimone and Lavington et al

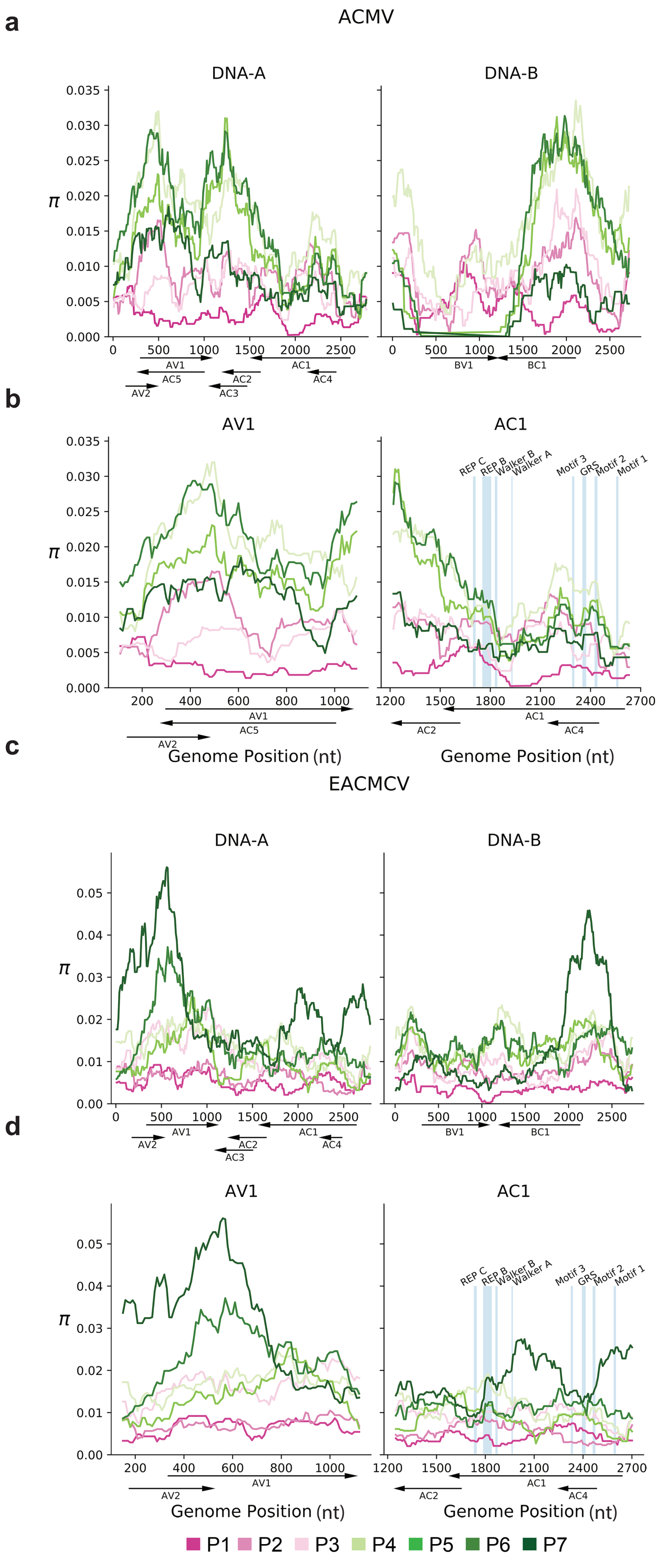
